# Gut Microbiota Mediates the Association between Diet Quality and Ectopic Adiposity: The Multiethnic Cohort Adiposity Phenotype Study

**DOI:** 10.64898/2026.04.10.717245

**Authors:** Songran Wang, Meredith A. Hullar, Keith Curtis, Sandi Kwee, Song-Yi Park, Christoph Rettenmeier, Kristine R. Monroe, Thomas Ernst, John Shepard, Lynne R. Wilkens, Loic Le Marchand, Johanna W. Lampe, Unhee Lim, Timothy W. Randolph

## Abstract

**Background:** Higher-quality diets have been associated with lower levels of ectopic fat deposited in the viscera and liver, which is hypothesized to be mediated in part by the gut microbiota.

**Objectives:** We tested this hypothesis in a multi-ethnic imaging study using global (microbiome-wide) testing as well as a high-dimensional multiple-mediators regression framework to identify bacterial genera in the human gut that mediate the association between diet quality and ectopic adiposity.

**Methods:** We analyzed the cross-sectional data of 1,400 older adults (age 60-77) from five racial/ethnic groups in the Multiethnic Cohort Adiposity Phenotype Study (2013-2016). Overall diet quality was defined by adherence to the MIND diet. The relative abundance of 151 bacterial genera was quantified from 16S rRNA gene sequencing of the stool samples. Visceral fat, liver fat, and the presence of MASLD (metabolic dysfunction-associated steatotic liver disease) were determined based on magnetic resonance imaging (MRI). We used high-dimensional mediation analysis (HDMA) to estimate gut microbial mediation in the linear regression of visceral fat or liver fat, or in logistic regression of MASLD, on the MIND adherence score, adjusted for potential confounders.

**Results:** Higher diet quality was associated with lower ectopic adiposity: 12% less visceral fat area, 23% less liver fat, and a 49% less likelihood of having MASLD, comparing the highest to the lowest quartile of the MIND score. Using a distance-based global test, we confirmed overall significant microbial mediation of the inverse diet-ectopic fat association. From HDMA, four bacterial genera were identified as mediating the protective association with visceral fat, with the largest mediation conferred by *Lachnospiraceae UCG.001* (12.2%). Two genera (*Lachnoclostridium*, *Weissella*) were shown to mediate the MIND association with both liver fat and MASLD. In particular, *Lachnoclostridium* mediated 13.6% of the liver fat association and 10.8% of the MASLD association, and *Lachnospiraceae UCG.001* additionally mediated 12.1% of the liver fat association.

**Conclusions:** Our results support the hypothesis that the gut microbiota contributes to conveying the effect of diet quality on preferred body fat distribution, e.g., involving bacteria that are known to produce short-chain fatty acids (Lachnospiraceae) or secondary bile acids (Lachnoclostridium).

## Introduction

Body fat distribution has long been recognized as more important than total fat mass for the risk of chronic diseases and premature deaths. ^57^ In prospective cohort studies, central or abdominal obesity, measured by waist circumference or its ratio to height or hip circumference, has shown stronger and more dose-response associations with morbidity and mortality than overall obesity, as approximated by body mass index (BMI), and remained associated after adjustment for BMI. ^13;17^ Beyond abdominal adiposity, the amount of ectopic fat around or inside organs, such as intra-abdominal visceral fat and hepatic fat, has shown a stronger relationship than abdominal subcutaneous fat to clinical biomarkers and metabolic dysfunctions.^37;48;56^ Visceral fat and liver fat, quantified accurately by magnetic resonance imaging (MRI), were each associated with the metabolic syndrome independently of MRI-measured abdominal subcutaneous fat or dual energy X-ray absorptiometry (DXA)-quantified total fat. ^44^

While visceral fat can be reduced in weight loss trials employing dietary energy restriction, exercise, or pharmaceutical agents^47;61^, we and others have observed that, even without change in weight or total body fat mass, high-quality dietary patterns may improve body fat distribution and lower the level of proportional visceral fat and hepatic fat. ^46;51;73;77^ For example, individuals scoring higher on a priori diet quality indices for adherence to healthy diets, or individuals improving on the score over time, were observed to have lower levels of visceral fat and liver fat, compared with their counterparts.^51^ Among the many diet quality indices that have been developed, ^71^ the Mediterranean diet, the Dietary Approach to Stop Hypertension (DASH) diet, and the Healthy Eating Index (HEI) have been frequently studied in the U.S. and fairly consistently associated with lower levels of visceral and hepatic adiposity. ^73;85^ Also, randomized controlled trials have shown that Mediterranean-pattern diets may outperform their isocaloric and healthy comparator diets in reducing visceral fat and other ectopic fat depots. ^27;83^

There may be multiple mechanisms underlying the relationship between diet quality and ectopic adiposity. High-quality diets have been linked to healthier lipid profiles, insulin sensitivity, and low systemic inflammation, which help sustain the lipogenic capacity of the subcutaneous adipose tissue as storage and prevent compensatory expansion of the visceral adipose tissue.^38^ Higher quality diets are also rich in substrates metabolized by gut bacteria, such as dietary fibers, resistant starches, and polyphenols, which help exert metabolic and endocrine effects for optimal regulation of adipose depots. ^81^ Notably, an investigation of blood metabolomics in relation to a visceral fat-associated diet identified two microbial metabolites as the leading correlates –– lower hippurate (a gut microbial metabolite of polyphenol-rich plant foods ^74^) and higher butyrylcarnitine (an intermediary product in gut microbial metabolism of L-carnitine to trimethylamine N-oxide) — and additionally detected the abundance of *Erysipelotrichaceae dolichum* to be associated with the visceral fat-associated diet. ^60^ Previous work by our group suggests that a positive association between pro-inflammatory diet and ectopic fat is mediated through specific bacterial genera such as *Fusobacterium* and *Lachnoclostridium*, bacteria known to induce host inflammation. ^46^

Given the emerging evidence that high-quality diets may suppress ectopic fat deposition in part through gut microbial composition and metabolism, we performed analyses to evaluate whether the effect of diet on ectopic fat is mediated by the host’s gut microbial composition. Our analysis was carried out in a cross-sectional MRI study that included participants from both sexes and five racial and ethnic groups. In this study, we previously observed a wide range of dietary patterns ^8^ and ectopic adiposity levels, ^44^ and also observed strong gut microbial associations with ectopic fat distributions. ^31;46^ In the current mediation analysis, we mainly examined the MIND diet, a hybrid between the Mediterranean and the DASH diets, that has been associated with lower risk of dementia and chronic diseases, ^54;55^ while also testing associations for the HEI diet. ^42;66^

Our primary focus is to evaluate the mediating effect of microbial abundances on the association between diet quality and ectopic adiposity using a multiple-mediators approach. In this framework, rather than estimating the marginal mediation effect of each genus separately, the analysis estimates the effect of each genus, conditional on (adjusting for) all others, within a penalized regression model that sparsely selects those with the strongest effect. ^84^ We implemented the high-dimensional mediation analysis (HDMA) of Gao et al. ^26^ as it provides de-biased estimates and inference for the genus-specific mediating effects ^15^ and allows for correlated mediators, which can influence the estimated mediation effect. ^64^ In addition, we tested for a global (microbiome-wide) mediation effect using the distance-based ‘MedTest’. ^88^ We acknowledge that by using a cross-sectional design and not controlling for all unmeasured confounders, a strictly causal interpretation may be compromised, but the results provide valuable insight on the potential role played by the microbiome in the relationship between diet and ectopic fat.

## Methods

### Study Population

We analyzed the cross-sectional data from the Multiethnic Cohort-Adiposity Phenotype Study (MEC-APS). The MEC-APS consists of 938 women and 923 men who have participated in the MEC since enrollment in 1993-1996^36^ and who underwent a detailed body composition study in 2013-2016 at age 60-77. ^44^ The APS participants were recruited from all five main racial and ethnic groups in the MEC based on self-report, including African American, Japanese American, Latino, Native Hawaiian, and White groups. The exclusion criteria were applied to known chronic viral hepatitis and diabetes treated with insulin, as well as recent smoking (*<*2 years) and any advanced disease conditions (e.g., advanced renal disease and thyroid conditions requiring medications), that might have altered the body fat distribution or the body fat depots’ association with biomarkers in our multi-omics investigation. ^44^ To ensure comparability across the sex-racial/ethnic groups that are known to have substantially different mean BMI levels and abdominal adiposity patterns, we recruited eligible volunteers into 60 strata as evenly as feasible by sex, five racial and ethnic groups, and six reported BMI levels (18.5-22.5, 22.5-25, 25-27, 27-30, 30-35 and 35-40 kg/m^2^). This led to oversampling heavier-weight Japanese Americans and lighter-weight African Americans and Native Hawaiians compared with their frequencies in the MEC and the general population, which served the intended purpose to robustly compare body fat distribution across the racial/ethnic groups, but the study design requires adjustment of total adiposity (BMI or total body fat mass) in the analyses. The study protocol was approved by institutional review boards at the University of Hawaii (UH) and the University of Southern California (USC), where the MEC participants were recruited and followed, including for the APS. All APS participants gave their informed consent, following the U.S. Common Rule on human subjects research. ^44^

### Data and Specimen Collections

The APS participants visited affiliated research clinics at UH in Honolulu, Hawaii, or USC in Los Angeles, California, and underwent anthropometric measurements (height, weight, waist and hip circumferences), whole-body DXA and abdominal MRI scans, collections of a fasting blood sample and a stool sample, following our established stool sample collection protocol, ^25^ and completion of study questionnaires, including the MEC questionnaire that collected demographic data and health and lifestyle history. ^44^

### Assessment of Dietary Intake and Overall Diet Quality

We re-administered the MEC quantitative food frequency questionnaire (QFFQ) at the time of the APS to assess the usual diet of the participants over the past year. ^36^ The QFFQ, composed of queries on over 180 food items/groups, was developed based on 3-day measured food records collected from the multiple racial/ethnic groups targeted in the MEC, and was validated for satisfactory correlations against three unannounced 24-hour recalls. ^70;79^ The overall quality of the diet was determined by computing the score for several established dietary pattern indices, based on the reported intake of component foods for each index. ^51^ We focused the current analysis on two specific indices. The MIND (the Mediterranean-DASH Intervention for Neurodegenerative Delay) integrates the Mediterranean diet and the DASH diet ^54;55^: for the MEC QFFQ, the original MIND score algorithm was modified to assess 9 of the 10 brain-healthy foods (excluding berries that were not asked directly in the MEC QFFQ) and all 5 unhealthy foods, for a scoring range of 0 to 14. ^62^ The HEI-2015 evaluates adherence to the 2015-2020 Dietary Guidelines for Americans (assessing 13 component foods for a scoring range of 0 to 100) based on a study population-independent scoring system. ^65^ We selected these two indices because they represent broadly applicable healthy diets and because they are well correlated with other similar indices. ^41;62;71^ For example, the Pearson correlation between the MIND and HEI-2015 scores was 0.65 overall, 0.69 in men, and 0.62 in women.

### Microbial Bioinformatic Processing

The stool samples were thawed and homogenized, and two of the aliquots were used to extract DNA as previously reported. ^31^ Total genomic DNA was quantified using the Qubit 1x dsDNA HS Assay Kit (Life Tech, Eugene, OR), and aliquots from the beginning and end of each period were shipped to MRDNA (Shallowater, TX) for library preparation and V1-V3 16S rRNA sequencing on a MiSeq (Illumina, San Diego, CA) following the manufacturer’s recommendations as previously reported. Sequence data was returned to the Fred Hutchinson Cancer Center (FHCC) using a secure server for subsequent bioinformatic analysis. Data was processed in USEARCH-UNOISE3 (usearch v11.0.667 i86linux64) ^22;23^ and QIIME2^7^. Amplicon single nucleotide variants (ASVs) were filtered by abundance to retain those *>* 0.001%. The taxonomy was assigned using the classify-sklearn module in QIIME2 with the SILVA 138 taxonomy using the V4 region. Phylogeny was constructed using the align-to-tree-mafft-fasttree command in QIIME2. This used MAFFT^34^ and FastTree2.

Samples that had fewer than 8656 sequence counts were removed from the dataset. This resulted in 1396 samples and 197 genera (from 12 phyla) detected with confidence. Finally, we filtered out genera that were present in fewer than 20% of the samples, resulting in a dataset with 151 genera. For statistical modeling, the centered log-ratio (CLR) was calculated on the genera taxon level to account for the compositional nature of the microbial abundances, ^2;28^ and account for the different number of total sequences collected in each sample. Briefly, the CLR transforms the abundance profile within each sample taking the log and subtracting of the sample’s mean. We note that the traditional CLR works when all values of the composition are positive, but it interrupts the ranking when zeros are handled separately. To preserve the overall ranking, we adopted a modified CLR which preserves zero abundance as a minimum value for each genus. ^82^

### Ectopic Fat Quantification and MASLD Classification

An abdominal MRI scan was acquired on 3-Tesla scanners to quantify the average visceral adipose tissue area (cm^2^) at four intervertebral segments between L1 and L5 and liver fat as a percentage in volume. ^44^ Visceral fat was scanned using an axial gradient-echo sequence with breath holds. ^43^ Liver fat was screened using a series of Dixon-type axial triple gradient-echo scans. ^29^ The presence of MASLD was determined based on the level of liver fat on imaging (*>* 5%), the amount of alcohol intake (men *<* 30g/day, women *<* 20g/day) to rule out alcoholic steatosis, and the presence of metabolic conditions, following the consensus definition. ^68^ The component metabolic conditions were assessed based on longitudinal reports of medical history, anthropometry (weight, height, waist circumference), clinical exams (blood pressure), blood biomarkers (glucose, insulin, triglycerides, high-density lipoprotein cholesterol), and an inventory of prescription medications.^37;44^

### Statistical Methods

#### Testing for Microbiome-wide Mediating Effects

We tested whether microbioime-wide profiles of abundance have a mediating effect on the association between diet quality and ectopic fat. For each combination of outcome *Y* (visceral fat, liver fat, and MASLD) and dietary exposure *X* (MIND and HEI), we performed a distance-based mediation test, MedTest, ^88^ with respect to the Bray–Curtis distance ^9^ between each pair of participants’ genus abundance profiles. We tested this using all 151 genera, as well as a subset of 36 genera which had previously been observed as associated with liver fat (10 positively 26 negatively) in this cohort. ^31^ The previous analysis modeled the association between liver fat and each genus, marginally, using beta-binomial regression adjusted for sex and percent total fat. The analysis here expands the analysis with a global test for the combined role of these genera in mediating the association between diet and fat.

#### Estimating Genus-Specific Mediating Effects Using Penalized Multiple-Mediators Regression

In view of the relatively large set of mediators, their potential collinearity, and the advantages of multiple-mediator modeling, ^64^ we adopted a regularized inference framework based on the de-sparsified LASSO^76;86^ as developed in the HDMA approach of Gao et al. ^26^ High-dimensional estimation is embedded within a mediation model designed to account for the large number of mediators and their potential correlation. This HDMA framework modifies the method by Zhang et al., ^87^ by implementing a simultaneous inference in a high-dimensional sparse regression model using a de-biased (de-sparsified) estimate.^86^ In our analysis, we estimated the indirect effect of a set of mediators *M* = (*M*_1_*, M*_2_*, … , M_p_*)^⊤^ (genera in the gut microbiome) in the pathway linking an exposure variable *X* (diet quality) to an outcome variable *Y* (ectopic fat). The outcome variable *Y* may be continuous or, as in the case of MASLD, binary. All mediation models were adjusted for key covariates, including age, sex, race/ethnicity, BMI (or total body fat mass), total calories, and alcohol intake. This multiple-mediator model is illustrated in Figure 1 where the effect of *X* on *Y* is transmitted both directly and indirectly through many potential mediators 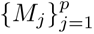.

**Figure 1:**
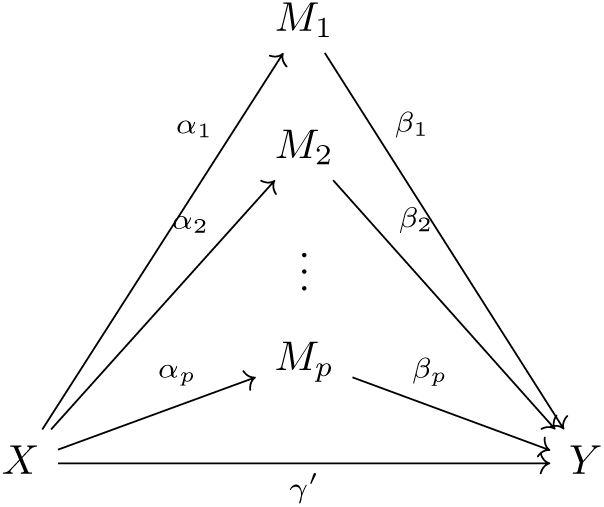
Multiple-mediators model. The potential mediating effects from all genera, 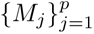, are modeled in a single framework. Here, *γ* represents the *total effect* between ectopic fat *Y* and diet quality *X*; *α_j_* is the association of diet *X* with genus *M_j_*; *β_j_* represents the effect of genus *M_j_* on fat *Y* after adjusting for diet; *γ*^′^ is the *direct effect* as adjusted for microbial abundances. The *indirect effect* is *α_j_β_j_* which, when scaled by the total effect *γ*, represents the *mediated effect*, *α_j_β_j_/γ*, by *M_j_*.

This is formally modeled as:

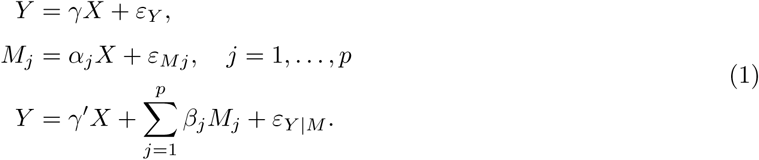

In this model, *α_j_*represents the effect of the exposure *X* on mediator *M_j_*, while *β_j_* denotes the effect of *M_j_* on the outcome *Y* after adjusting for *X*. The coefficient *γ* captures the total effect of *X* on *Y* , and *γ*^′^ represents the direct effect after accounting for the mediators. The product *α_j_β_j_* quantifies the indirect effect of *X* on *Y* through mediator *M_j_*, and the difference *γ* − *γ*^′^ corresponds to the overall mediation effect aggregated across all mediators. The proportion of exposure-on-outcome that is mediated by *M_j_*, denoted by (*α_j_β_j_*) */γ* × 100, represents the extent to which the mediator has an effect on the total association. We tested the null hypothesis *H*_0_ : *α_j_β_j_* = 0, implying no mediation effect via *M_j_*, using a joint significance approach. A mediator was declared significant only if both *α_j_* and *β_j_* were significantly different from zero at the pre-specified threshold *p <* 0.05. As outlined next, the HDMA testing framework conducts simultaneous inference with respect to a sparse mediation-vector estimate based on the de-biased LASSO estimator, without a post-hoc adjustment for multiple comparisons. ^84;86^ The framework allows for covariate adjustment and we included the covariates of age, sex, race/ethnicity, BMI (or total body fat mass), total calories, and alcohol intake.

##### The HDMA Estimation and Inference Procedure

Our goal was to estimate the indirect effect of a set of variables *M* = (*M*_1_*, M*_2_*, … , M_p_*)^⊤^ for their potential mediating effect on the association between *X* (diet quality) and *Y* (ectopic fat). As described above, HDMA high-dimensional sparse regression with respect to a de-biased estimation procedure and the inference reported arises from a single, high-dimensional model accounting for the fact that multiple predictors contribute to a potential mediation effect. The HDMA proceeds in three steps. The first step applies Sure Independent Screening (SIS) ^24^ to reduce the number of mediators from *k* to a moderate dimension *d*, calculated as *d* = ⌈*n/* log(*n*)⌉, for continuous outcomes, or *d* = ⌈*n/*(2 log(*n*))⌉, for binary outcomes, where *n* is the number of observations. The screening is based on marginal associations of each *M_j_* regressed on *Y* , for continuous *Y* , or on *X* for binary *Y*. Mediators with the strongest associations are retained for further analysis. In our setting, although the sample size exceeds the number of genera, the set of potentially mediating genera is still relatively large and may have strong correlations among them and thus the SIS procedure assures that the least relevant genera are excluded.

The second step performs high-dimensional inference on the retained set of *d* mediators. Specifically, the following model is fit:

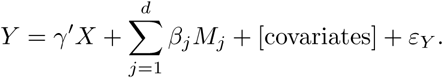

Although the number of mediators in this regression model is moderate (*d < n* in our analysis), the use of the de-biased LASSO estimate in this step selects for mediating genera while correcting for the estimation bias introduced by penalization and enables valid asymptotic inference on each *β_j_*. For each *j* ∈ {1*, … , d*}, a two-sided *p*-value is computed for testing *H*_0_ : *β_j_* = 0, and only those mediators with *p_j_ <* 0.05 are retained for the next step of mediation testing.

In the third step, for the subset of mediators passing the significance threshold on the *β_j_* coefficients, the corresponding coefficients *α_j_* are estimated. For continuous *M_j_*, this is done by fitting linear models of the form *M_j_* = *α_j_X*+[covariates]. The indirect effect was computed as the product *α_j_β_j_*, and the joint significance test for mediation was performed by taking the maximum of the *p*-values for *α_j_* and *β_j_* , denoted 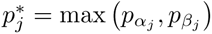. A mediation effect was declared significant if 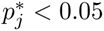.

To estimate the total effect *γ*, we fit a standard regression model of *Y* ∼ *X*+[covariates] and extracted the coefficient of *X*. Using the estimated indirect effect and total effect, we calculated the proportion of the total effect mediated by each selected mediator.

##### Reported Quantities

For each identified mediator exhibiting a significant mediation effect, we reported the following statistical quantities: the estimates for *α_j_, β_j_*, and *γ*; the estimated indirect effect *α_j_β_j_*; the proportion of mediation (*α_j_β_j_*) */γ*× 100; and the joint p-value for the mediation effect 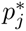. These estimates provided comprehensive insights into both the strength and statistical significance of each mediation pathway.

## Results

### Descriptive Characteristics of the Study Population

Table 1 shows the characteristics of the APS participants, overall and by quartile of the MIND diet score. The age and sex distributions were similar across the levels of diet quality. Groups with a higher MIND score tended to have more White participants and fewer Latino or Native Hawaiian participants. Individuals in the higher MIND score categories were also likely to have higher education and a lower history of hypertension or diabetes. As expected, the mean score for the HEI diet (HEI-2015) also increased with increasing levels of the MIND diet score, as were caloric intake and alcohol consumption. Also, adherence to the recommended physical activity was higher, and anthropometric measurements (BMI, waist circumference, visceral fat area, liver fat, and percentage of MASLD) were lower in higher MIND score categories.

**Table 1:** Description of the APS participants overall and by the MIND score quartile.

| | Overall | MIND Quartile 1<br>($\leq 6.5$) | MIND Quartile 2<br>($6.6 \leq 7.5$) | MIND Quartile 3<br>($7.6 \leq 9.0$) | MIND Quartile 4<br>($> 9.0$) |
| --- | --- | --- | --- | --- | --- |
| No. participants | 1396 | 388 | 306 | 443 | 259 |
| Age, y (SD) | 69.2 (2.8) | 69.2 (2.9) | 69.5 (2.7) | 69.1 (2.7) | 69.0 (2.7) |
| Female % | 53 | 51 | 52 | 55 | 56 |
| **Race and Ethnicity, %** |  |  |  |  |  |
| White | 21 | 14 | 21 | 23 | 28 |
| African American | 16 | 18 | 17 | 16 | 16 |
| Latino | 21 | 22 | 22 | 21 | 17 |
| Asian/Japanese American | 26 | 28 | 25 | 25 | 25 |
| Native Hawaiian | 16 | 19 | 15 | 15 | 14 |
| Education (>12th grade) | 81 | 79 | 76 | 83 | 85 |
| **Cardiometabolic history, %** |  |  |  |  |  |
| Hypertension | 49 | 54 | 54 | 47 | 39 |
| Diabetes | 16 | 18 | 18 | 16 | 9 |
| Smoking (past) | 37 | 39 | 38 | 34 | 37 |
| **Lifestyle and anthropometry** |  |  |  |  |  |
| HEI-2015, score (SD) | 71.6 (10.4) | 63.2 (8.9) | 69.7 (8.8) | 74.6 (8.0) | 81.2 (6.7) |
| Alcohol intake, g/day (SD) | 4.0 (6.4) | 3.2 (6.3) | 3.7 (6.1) | 4.3 (6.4) | 5.1 (6.9) |
| Caloric intake, kcal/day (SD) | 1819 (859) | 1822 (913) | 1736 (817) | 1753 (821) | 2025 (857) |
| Physical activity (>150 min/week), % | 86 | 79 | 90 | 87 | 93 |
| Body mass index, kg/m<sup>2</sup> (SD) | 27.9 (4.8) | 29.2 (4.9) | 28.3 (4.6) | 27.4 (4.7) | 26.1 (4.5) |
| Waist circumference, cm (SD) | 95.3 (12.2) | 98.5 (11.8) | 96.7 (11.9) | 94.2 (12.3) | 90.4 (11.4) |
| Visceral fat area, cm<sup>2</sup> (SD) | 166 (83) | 188 (85) | 178 (82) | 159 (82) | 129 (71) |
| Liver fat, mean% (SD) | 5.7 (4.6) | 6.7 (5.0) | 5.8 (4.5) | 5.6 (4.6) | 4.1 (3.5) |
| MASLD, mean% | 38 | 48 | 41 | 39 | 19 |

### Association between Diet Quality and Ectopic Fat

As with other diet quality indices, ^51^ the MIND diet score was inversely associated with visceral fat, liver fat, and the presence of MASLD even after adjusting for BMI and total caloric intake (Supplementary Table S1). The mean visceral fat area was 12% lower, while the mean liver fat was 23% lower, among the participants in the highest quartile for the MIND diet score compared with those in the lowest quartile. Similarly, those in the highest quartile for the MIND score were 61% less likely to have MASLD. The similar inverse relationship between adherence to the HEI diet and ectopic fat phenotypes in the MEC-APS was previously presented. ^51^ The diet-ectopic fat associations for either the MIND or HEI diet and for any of the visceral or hepatic adiposity outcomes were generally stronger among women than among men (Supplementary Table S1).

### Gut Microbial Mediation of the Association between Diet Quality and Ectopic Fat

#### Global testing for mediation

We first applied ‘MedTest’, the distance-based approach of Zhang et al., ^88^ to test for the mediating effect of all microbial abundances, together, in the association between each dietary index and each type of ectopic fat. This test is analogous to PERMANOVA, ^4^ a global test commonly applied in microbiome analysis. Using the Bray-Curtis distance between pairs of samples (each sample is a vector of 151 genus abundances), we observed strong evidence that the gut microbiome mediates the various associations between diet and fat. Indeed, for each combination of outcome *Y* (visceral fat, liver fat, and MASLD) and dietary exposure *X* (MIND and HEI), the MedTest for a microbiome-wide mediating effect produced p-values smaller than 0.0001.

We also considered a subset of genera that were previously reported as marginally associated with liver fat within this same cohort of MEC participants. ^31^ Of these, 10 were positively associated and 26 negatively associated (see Supplemenatary Table S2). Using this subset of 36 genus abundances in the distance-based MedTest, we observed mediating effects involving the MIND diet and all three types of ectopic fat: liver fat (p=0.001), visceral fat (p=0.002), MASLD (p=0.001). We note that the updated bioinformatic processing used in the present analysis identified a slightly different set of taxa, leading to incomplete overlap in the genera studied here versus previously.

#### High-dimensional mediation analysis

We investigated the gut microbiome’s mediating effect on the inverse association between the MIND diet and visceral fat levels (shown in Supplementary Table S1) using 151 genera in the regularized regression framework of HDMA. The genera having the most prominent mediating effect on the diet quality-ectopic fat relationship are shown in Tables 2–4.

**Table 2:**
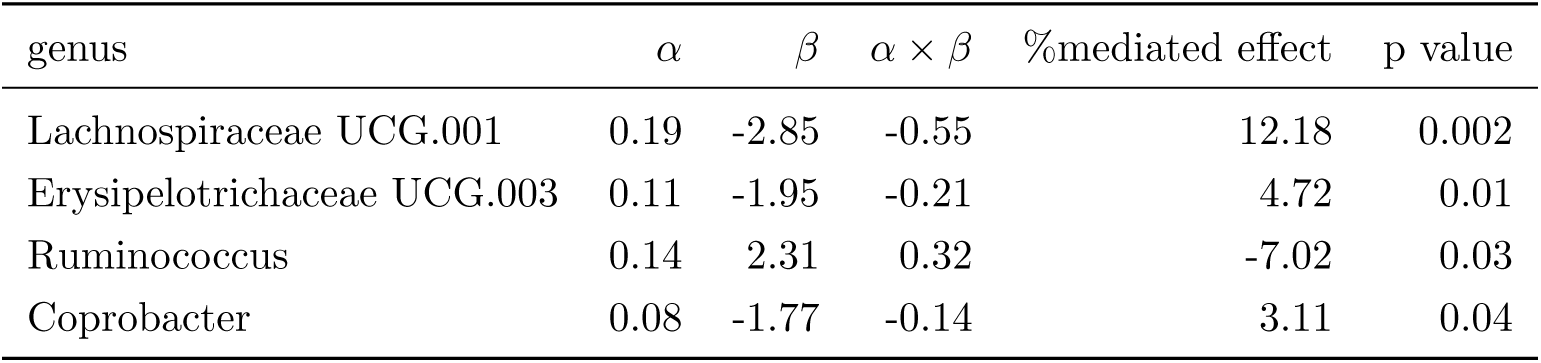
Gut microbial mediators of the association between the MIND diet *X* and visceral fat *Y* , in the HDMA model (1) with respect to 151 genera adjusted for sex, race/ethnicity, age, BMI, total calories, and alcohol intake. The estimated total effect was *γ* = −4.51.

**Table 3:**
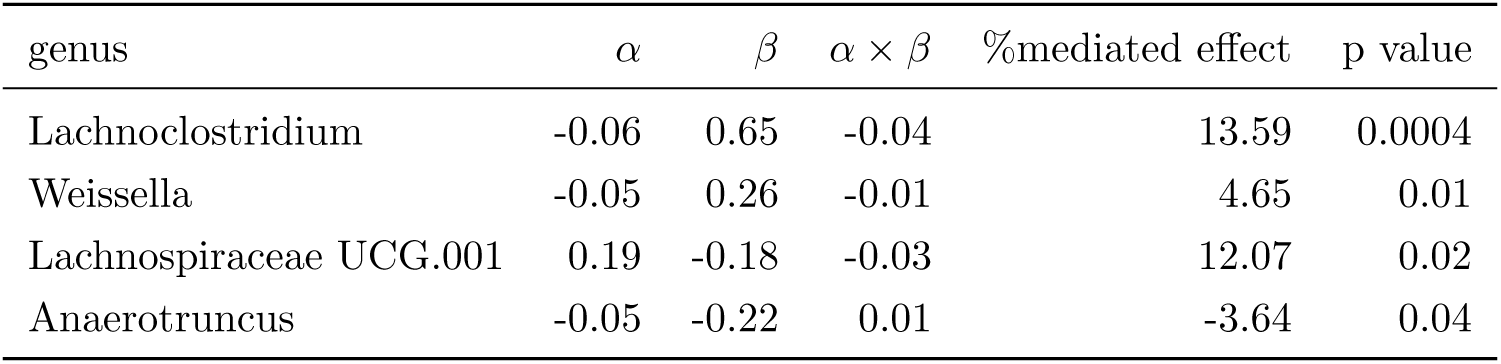
Gut microbial mediators of the association between the MIND diet *X* and liver fat *Y* , in the HDMA model (1) with respect to 151 genera adjusted for sex, race/ethnicity, age, BMI, total calories, and alcohol intake. The estimated total effect was *γ* = −0.28.

**Table 4:**
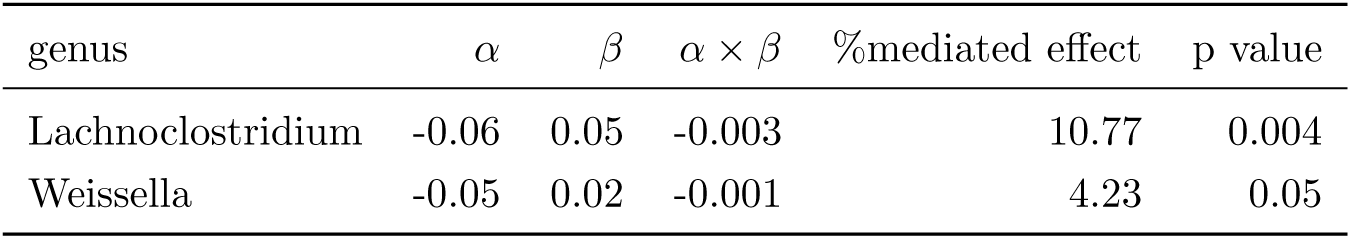
Gut microbial mediators of the association between the MIND diet *X* and MASLD *Y* , in the HDMA model (1) with respect to 151 genera adjusted for sex, race/ethnicity, age, BMI, total calories, and alcohol intake. The estimated total effect was *γ*= −0.03.

The total effect of the MIND diet on visceral fat levels was mediated by *Lachnospiraceae UCG.001*, *Erysipelotrichaccae UCG.003*, *Ruminococcus*, and *Coprobacter* at 12.2%, 4.7%, 7.0%, and 3.1%, respectively (Table 2). *Ruminococcus* had a negative mediating effect (i.e., diminishing the protective effect of the MIND diet against visceral adiposity), while the mediating effect of the others appeared to support the visceral fat-lowering effect of the MIND diet. Specifically, *Ruminococcus* was positively associated with both the MIND diet (*α* = 0.14) and visceral fat (*β* = 2.31), suggesting that the inverse MIND-visceral fat association may be attenutated by the presence of *Ruminococcus*.

Several genera were identified as significant mediators in the association between the MIND diet and liver fat levels, including *Lachnoclostridium* (13.6% mediation), *Weissella* (4.6% mediation), *Lachnospiraceae UCG.001* (12.1% mediation), and *Anaerotruncus* (−3.6% mediation) (Table 3). Of these, *Anaerotruncus* had a negative mediating effect, due to its correlation with lower MIND scores and also lower liver fat.

Notably, the top two genera for the MIND-liver fat mediation, *Lachnoclostridium* (10.8% mediation) and *Weissella* (4.2% mediation), were also prominent in mediating the MIND-MASLD association (Table 4).

It should be noted that the scales for these ectopic fat quantities differ, so the total effect sizes of the diet-ectopic fat associations, *γ*, in Tables 2–4 are not directly comparable to each other. However, the percentage of the effect mediated by the microbiome can be compared across these analyses.

Similar associations and mediating effects are observed when the analysis is stratified by sex; see Supplementary Tables S3–S5. Our analysis of another diet quality index, the HEI diet score, with ectopic fat phenotypes yielded similar lists of mediating gut bacteria. These are shown in Supplementary Tables S6–S8.

## Discussion

In this cross-sectional study, we evaluated the role of human gut microbiota in the ectopic fat-lowering effect of the MIND diet using global (distance-based) testing and high-dimensional mediation analysis. The results point to a significant role played by the microbiome in partially mediating the inverse relationship between diet quality and ectopic adiposity, while accounting for potential confounding by sex, race/ethnicity, age, and total caloric and alcohol intakes.

We identified several gut microbial genera, four each for visceral fat and hepatic fat (both liver fat and MASLD), that mediated the ectopic fat-lowering effect of the MIND diet. For visceral fat, *Lachnospiraceae UCG.001*, *Erysipelotrichaceae UCG.003*, and *Coprobacter* had a positive mediating effect of up to 12.2% of the total dietary effect, while *Ruminococcus* had negative mediating effect. *Lachnospiraceae UCG.001* had a similarly strong mediating effect (12.1% of the total effect) for liver fat, closely following the mediation by *Lachnoclostridium* (13.6% of the total effect). *Lachnoclostridium* and *Weissella* were observed to mediate both the continuous and dichotomous outcomes for hepatic steatosis. *Anaerotruncus* was additionally identified as a mediator for liver fat. These mediation effects for ectopic fat depots were independent of total adiposity (BMI or total fat mass measured by DEXA). Generally, the findings were similar between women and men, as well as for the mediation of the HEI diet effect. These results are shown in Supplementary Tables S3–S8.

Our study was motivated by observations that overall diet quality may affect body fat distribution,^46;51;73;77^ which suggest that an increase in diet quality leads to a decrease in ectopic fat. Since these fat depots carry higher metabolic risks, it is important to understand the mechanisms underlying this relationship. As in our previous study, where we observed that diet quality based on adherence to the DASH, Mediterranean, or HEI diets was inversely related to ectopic fat, ^51^ the current analysis indicates that the MIND diet, a hybrid of the DASH and the Mediterranean diets, is associated with up to 12% less visceral fat, 23% less liver fat, and 61% lower likelihood of MASLD when comparing the highest to the lowest quartile of the diet score. The MIND and the HEI diets showed a similar magnitude of associations with ectopic fat, which was stronger than the association observed for the dietary inflammatory index (DII)^46^: 10% lower visceral fat, 12% lower liver fat, and 37% lower likelihood of MASLD when comparing the lowest to the highest quartile of DII (data otherwise not shown). High-quality diets, such as those rich in beneficial bioactive compounds (e.g., phytochemicals, monounsaturated fatty acids), have been suggested to increase fatty acid oxidation, inhibit de novo lipogenesis, and facilitate browning and thermogenesis of white adipose tissue. ^27;38^ These mechanisms may have a greater effect on the metabolically more dynamic ectopic fat depots than on subcutaneous fat, as evidenced in a randomized controlled trial comparing the efficacy of the high-quality Mediterranean diet and the low-fat (high-carbohydrate) diet on various body fat depots over 18 months.^27^ Thus, our goal was to test whether some of the effects of diet quality on ectopic fat are exerted by influencing gut microbial composition, as we previously found gut microbial mediation of the relationship between a proinflammatory diet and ectopic fat in a simpler mediation analysis. ^46^

Visceral fat accumulation has been separately associated with dietary patterns and with the role of gut microbiome in SCFA generation. We found contrasting associations in the mediation between diet and visceral fat for *Lachnoclostriduium* and *Ruminococcus*. *Lachnoclostridium* as inversely association with HEI and positively associated with visceral fat. An increase in *Lachnoclostridium* has been associated with high-fat intake and lower HEI.^5^ *Lachnoclostridium* produces acetate, a precursor in lipogenesis, and has been positively correlated with visceral fat^58^ and acetate has been shown to mediate ≈10% of the total effect of *Lachnoclostridium* on visceral fat. ^33^ *Lachnoclostridium* is also associated with secondary bile acid (SBA) production. ^80^ SBAs play pivotal roles in modulating host lipid, glucose, and energy homeostasis ^1;53^ through the activation of nuclear receptors-—farnesoid X receptor (FXR), pregnane X receptor, and vitamin D receptor—and a G protein-coupled receptor (TGR5).^16^ as well as through the regulation of glucagon-like peptide (GLP-1) synthesis. ^75;78^ In contrast, *Ruminococcus* and *Bacteroides* were positively associated with the HEI and MIND diets and negatively associated with visceral fat in our data. *Ruminococcus* and *Bacteroides* are associated with dietary patterns including long-term fruit and vegetable consumption ^18;19^ and are known acetate producers. ^49;52;67;69^ In diets high in plant fibers, cross-feeding of acetate to other members of the gut microbial community effectively reduces circulating levels of acetate available for ectopic fat metabolism and produces butyrate (via condensation of two acetates), a SCFA associated with reduced inflammation. ^21;58^

In recent years, advancements in the understanding of MASLD pathogenesis have led to a shift from the traditional “two-hit” hypothesis to the more comprehensive “multiple-hit” hypothesis, ^10;63^ which emphasizes the critical role of the gut–liver axis in the onset and progression of the disease. ^39;40^ Increasing evidence indicates that the gut microbiota and its metabolites are critically involved in modulating hepatic inflammation and oxidative damage. ^14;20;35;45^ In addition, dietary interventions have been recognized as an essential component in the prevention and management of MASLD by modulating metabolic risk factors and liver inflammation. ^85^ In our mediation analysis we found that *Lachnoclostridium*, which can convert choline to trimethylamine (TMA), explained the most variation in the association between diet and MASLD. *Lachnoclostridium* was negatively associated with the MIND and HEI indices which reflects other studies showing a positive association with diets high in red meat and fat consumption. Diets high in red meat are also high in choline which can be converted to TMA by *Lachnoclostridum* and then further oxidized to trimethylamine-N-oxide (TMAO), a metabolite that has been associated with the severity of MASLD-induced hepatic inflammation. ^11;19;72;89^ Furthermore, SBAs associated with diets higher in red meat and animal fats also are associated with increased MASLD.^59^ Several clinical studies have shown that the concentration of the SBA deoxycholate (DCA) is greater in patients with MASLD than in healthy volunteers. ^32^ MASLD patients exhibit elevated serum levels of bile acids, along with an increase in SBAs, and a higher abundance of bile acid-metabolizing gut bacteria. ^30^

While alcoholic liver disease is attributed to excessive alcohol consumption, endogenous ethanol production by microorganisms can be substantial and depends on the diet composition. ^50^ *Weisella* was inversely associated with the MIND diet and positively associated with MASLD explaining 4% of the total variation between diet and MASLD. This is likely because *Weisella* uses heterofermentation to produce ethanol and lactate. ^6^ Changes in ethanol levels produced by microorganisms have been described in the spectrum of MASLD stages, contributing to liver toxicity and progression toward liver fibrosis and chronic liver disease. ^3;12;50^

This study has several strengths. We demonstrated the microbial mediation of the diet quality on ectopic adiposity through several analytical approaches and consistent patterns of results. Since the number of genera of interest is relatively large, we applied methods that account for the entire microbial content, rather than modeling each genus in isolation. In particular, the MedTest quantifies the microbiome using sample-wise distances and hence tests for a microbiome-wide role in mediating the diet quality/ectopic fat relationship observed in these participants. Second, by using a multiple-mediators model as implemented within the HDMA framework, the interpretation of genus-specific mediation is conditional on all genera rather than modeling each one in isolation. This approach also leverages recent advances on high-dimensional inference, allowing for a rigorous and efficient analyses of multiple bacterial mediators. The study sample of generally healthy older adults was drawn from a population-based cohort representative of both sexes and five racial/ethnic groups, capturing a wider range of diet quality and ectopic fat levels. Dietary intake was assessed using a comprehensive multiethnic QFFQ, and the resulting diet quality indices have been associated with various health outcomes in the cohort, demonstrating the construct validity of the dietary assessment in the MEC. Rigorous assessment was applied to quantify visceral fat and liver fat by gold-standard MRI protocols, and to define MASLD based on objective biomarker tests and direct medication inventories.

A primary limitation of the study is that findings from cross-sectional studies are susceptible to bias in determining causal inferences for mediation. Although the direction of mediation estimated for the top mediator bacteria is consistent with their known functions, as discussed above, our results and interpretation are limited to structural composition as assessed with 16S rRNA sequencing. Also, a larger study might have allowed detection of less common gut microbial genera as significant mediators of diet quality.

In summary, our findings add to the growing evidence that the gut microbiota plays a critical role in conveying the effects of diets. Future metagenomic studies are warranted to investigate the microbial metabolic functions involved in mediation.

## Acknowledgment

We thank the study participants who generously donated their time and effort to the Multiethnic Cohort and the Adiposity Phenotype Study (“Body Imaging Study”) research. We also gratefully acknowledge the contributions of a large team of research staff who assisted with these studies.

## Authors’ Contributions/Responsibilities

The authors’ contributions were as follows: LLM led the APS study; LLM, LRW, UL, and KRM contributed to the design, recruitment, and data collection for the APS as relevant to the current analysis; MAH, JWL, and TWR generated the gut microbiome data; TE ascertained the MRI measurements, and TE and CR provided input on MR data analysis; SK provided clinical input on liver outcome analysis; TWR, MAH, and UL conceived the manuscript; SW and TWR performed the data analysis; SW, TWR, MAH, and UL drafted the manuscript; and all authors contributed to revisions of the manuscript and approved the final version submitted to the journal.

## Disclosure of Conflict of Interest

None of the authors reported a conflict of interest related to this study or their institutional responsibilities.

## Funding

This work was supported by grants from the National Cancer Institute (P01 CA168530, U01 CA164973, P30 CA071789, P30 CA015704) and the National Institute on Minority Health and Health Disparities (R01 MD018265) at the U.S. National Institutes of Health.

## Data Availability Statement

The MEC-APS data are available upon request from the Multiethnic Cohort Research Committee. The Data Sharing Policy for the MEC and ancillary studies is provided at: https://www.uhcancercenter.org/for-researchers/mec-data-sharing.

## Supplementary Material

**Figure S1:**
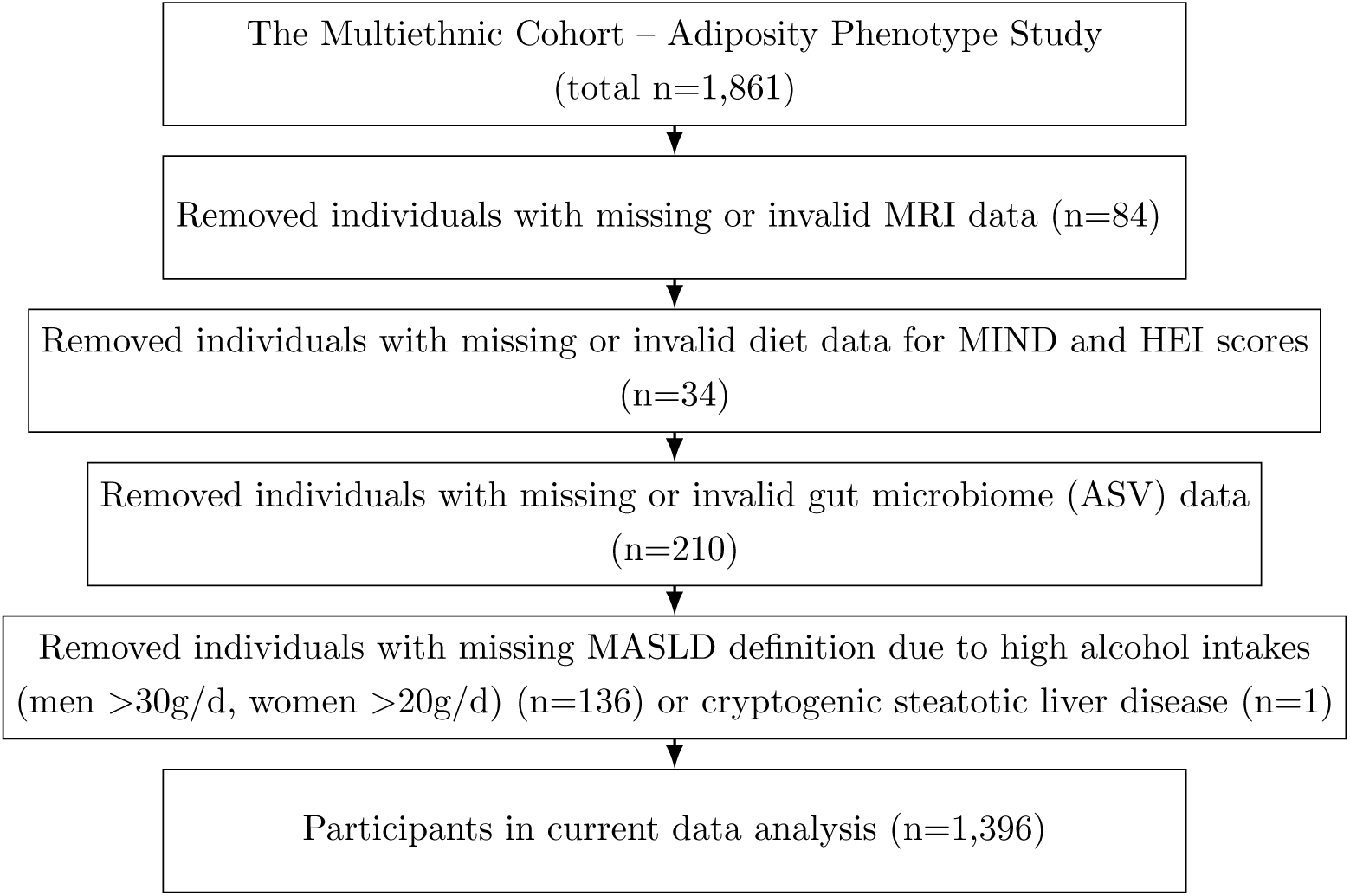
Flowchart for selection of study participant in the current analysis.

**Table S1:** Association between overall diet quality and ectopic fat phenotypes. The Pearson correlation between the MIND and HEI-2015 scores was 0.65 overall, 0.69 in men, and 0.62 in women.

|  | MIND |  |  |  |  | HEI |  |  |  |  |
| --- | --- | --- | --- | --- | --- | --- | --- | --- | --- | --- |
|  | Q1 | Q2 | Q3 | Q4 | p trend | Q1 | Q2 | Q3 | Q4 | p trend |
| **Visceral Fat** |  |  |  |  |  |  |  |  |  |  |
| Overall | 172 (167, 177) | 172 (167, 177) | 163 (158, 169) | 154 (149, 159) | <0.0001 | 174 (169, 180) | 168 (163, 173) | 163 (158, 168) | 156 (150, 161) | <0.0001 |
| Men | 200 (193, 208) | 201 (193, 210) | 198 (189, 206) | 183 (175, 191) | 0.005 | 203 (195, 211) | 195 (187, 203) | 195 (187, 204) | 187 (178, 195) | 0.07 |
| Women | 140 (133, 145) | 142 (136, 148) | 129 (123, 135) | 126 (121, 132) | 0.0002 | 145 (138, 151) | 140 (134, 146) | 132 (126, 138) | 122 (117, 128) | <0.0001 |
| **Liver Fat** |  |  |  |  |  |  |  |  |  |  |
| Overall | 6.2 (5.8, 6.6) | 5.7 (5.3, 6.2) | 5.5 (5.0, 5.9) | 4.7 (4.3, 5.1) | <0.0001 | 6.2 (5.8, 6.6) | 6.0 (5.6, 6.4) | 5.3 (4.9, 5.7) | 4.6 (4.2, 5.0) | <0.0001 |
| Men | 6.0 (5.5, 6.5) | 5.2 (4.5, 5.7) | 5.7 (5.1, 6.3) | 4.6 (4.0, 5.1) | 0.001 | 5.9 (5.3, 6.4) | 5.6 (5.1, 6.2) | 5.1 (4.5, 5.6) | 4.8 (4.2, 5.4) | 0.03 |
| Women | 6.3 (5.7, 6.9) | 6.3 (5.7, 6.9) | 5.3 (4.7, 5.9) | 4.9 (4.4, 5.5) | 0.0009 | 6.6 (5.9, 7.2) | 6.4 (5.8, 7.0) | 5.7 (5.1, 6.3) | 4.5 (3.9, 5.1) | <0.0001 |
| **MASLD** |  |  |  |  |  |  |  |  |  |  |
| Overall | 1.0 (Ref) | 0.82 (0.60, 1.13) | 0.89 (0.65, 1.22) | 0.51 (0.37, 0.70) | 0.0002 | 1.0 (Ref) | 0.88 (0.64, 1.20) | 0.85 (0.62, 1.16) | 0.52 (0.37, 0.72) | 0.001 |
| Men | 1.0 (Ref) | 0.76 (0.48, 1.20) | 1.08 (0.68, 1.70) | 0.58 (0.36, 0.92) | 0.047 | 1.0 (Ref) | 0.96 (0.61, 1.49) | 1.10 (0.70, 1.73) | 0.64 (0.39, 1.06) | 0.21 |
| Women | 1.0 (Ref) | 0.93 (0.59, 1.46) | 0.80 (0.51, 1.25) | 0.50 (0.32, 0.77) | 0.01 | 1.0 (Ref) | 0.82 (0.53, 1.29) | 0.71 (0.46, 1.12) | 0.44 (0.27, 0.69) | 0.003 |

**Table S2:**
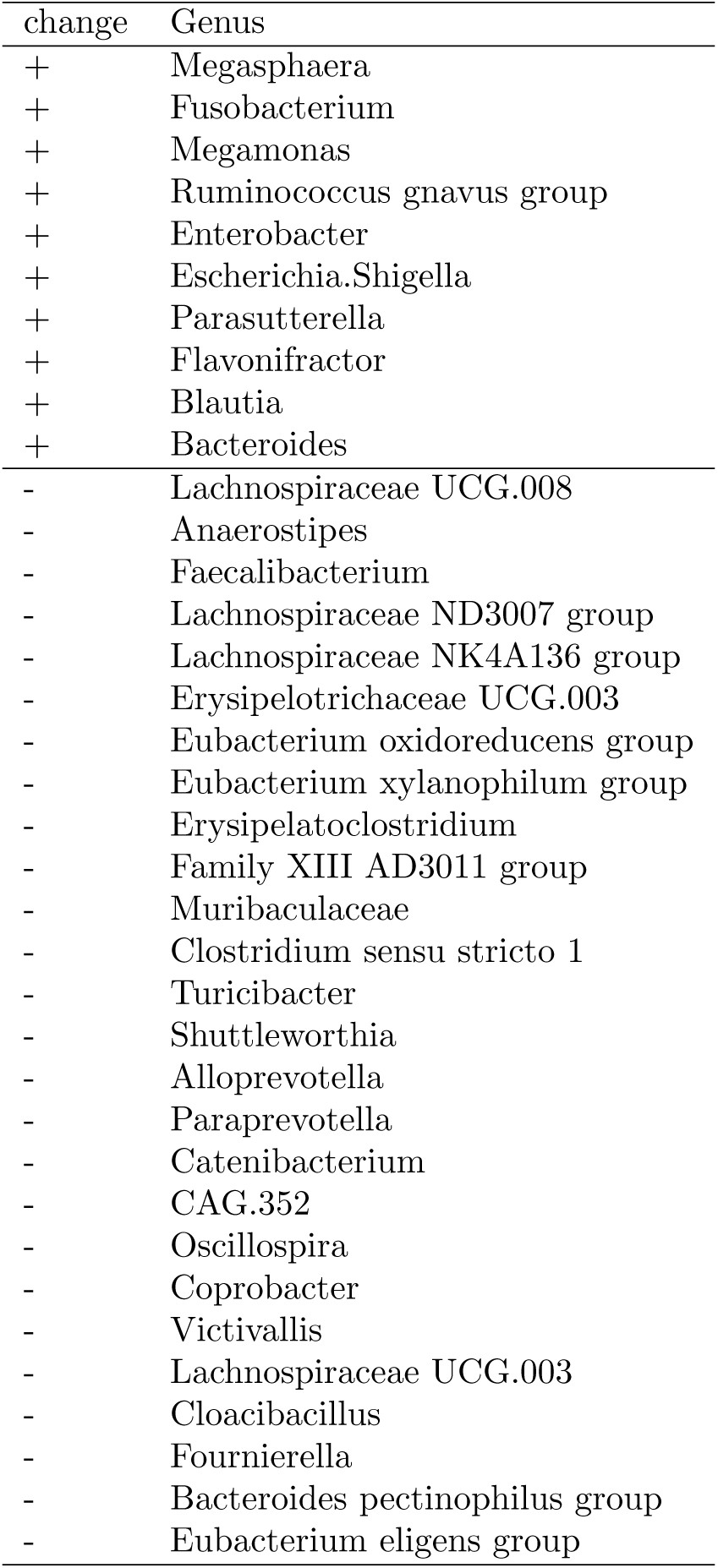
Genera whose abundance was previously observed to be increased (+) or decreased (−) among participants in the MEC cohort who exhibited high liver fat.

### Microbial mediators of the association between the MIND index and ectopic fat, stratified by sex

**Table S3:** Genera mediating an association between *X*=MIND and *Y* =visceral fat stratified by males and females.

| Group | genus | $\alpha$ | $\beta$ | $\alpha \times \beta$ | %mediated effect | p value |
| --- | --- | --- | --- | --- | --- | --- |
| Male<br>$\gamma = -3.86$ | Catenibacterium | -0.17 | 2.65 | -0.45 | 11.66 | 0.02 |
|  | Candidatus Stoquefichus | -0.14 | -3.85 | 0.55 | -14.29 | 0.02 |
| Female<br>$\gamma = -3.46$ | Lachnospiraceae UCG.001 | 0.19 | -2.84 | -0.53 | 15.26 | 0.005 |
|  | Eubacterium eligens group | 0.16 | 2.62 | 0.42 | -12.26 | 0.01 |
|  | Lachnoclostridium | -0.06 | 4.79 | -0.31 | 8.86 | 0.02 |
|  | Megasphaera | -0.10 | -2.23 | 0.23 | -6.75 | 0.03 |
|  | Eggerthella | -0.08 | -2.72 | 0.21 | -6.20 | 0.03 |
|  | Lactonifactor | -0.05 | -3.40 | 0.17 | -5.02 | 0.04 |

**Table S4:** Genera mediating an association between *X*=MIND and *Y* =liver fat stratified by males and females.

| Group | genus | $\alpha$ | $\beta$ | $\alpha \times \beta$ | %mediated effect | p value |
| --- | --- | --- | --- | --- | --- | --- |
| Male<br>$\gamma = -0.26$ | Incertae Sedis | -0.07 | -0.40 | 0.03 | -11.39 | 0.01 |
|  | Weissella | -0.10 | 0.30 | -0.03 | 11.20 | 0.03 |
|  | Lachnoclostridium | -0.05 | 0.57 | -0.03 | 10.68 | 0.04 |
|  | Clostridia UCG.014 | 0.18 | -0.17 | -0.03 | 11.96 | 0.05 |
|  | Streptococcus | -0.09 | -0.20 | 0.02 | -7.28 | 0.05 |
| Female<br>$\gamma = -0.25$ | Lachnoclostridium | -0.06 | 0.83 | -0.05 | 21.20 | 0.008 |
|  | Lachnospiraceae UCG.001 | 0.19 | -0.22 | -0.04 | 16.38 | 0.047 |

**Table S5:** Genera mediating an association between *X*=MIND and *Y* =MASLD stratified by males and females.

| Group | genus | $\alpha$ | $\beta$ | $\alpha \times \beta$ | %mediated effect | p value |
| --- | --- | --- | --- | --- | --- | --- |
| Male<br>$\gamma = -0.02$ | Parabacteroides | -0.12 | 0.04 | -0.005 | 26.47 | 0.01 |
|  | Faecalitalea | -0.18 | -0.02 | 0.004 | -26.97 | 0.04 |
|  | Lachnoclostridium | -0.05 | 0.06 | -0.003 | 17.17 | 0.04 |
| Female<br>$\gamma = -0.03$ | Colidextribacter | -0.07 | 0.04 | -0.003 | 11.16 | 0.02 |
|  | Lachnoclostridium | -0.06 | 0.05 | -0.003 | 10.74 | 0.03 |

### Gut microbial mediation of the association between the HEI score and ectopic fat

The genera with the strongest mediating effect between liver fat and HEI were *Lachnoclostridium* (13.6%) and *Lachnospiraceae UCG.001* (7.8%). Others include *Oscillibacter*, *Clostridia UCG.014*, *Weissella*, and *Bilophila*. Negative mediating effects were observed with *Collinsella*, *Aldercreutzia*, and *Anaerotruncus*.

Mediating the association between visceral fat and HEI were again *Lachnospiraceae UCG.001* (14.6%) and *Lachnoclostridium* (8.8%), along with other positive mediators of *Negativibacillus*, *Tyzzerella*, *Acidaminococcus*, and *Erysipelotrichaceae UCG.003*. Negative mediation was observed with *Ruminococcus* and *Adlercreutzia*.

As for the HEI-MASLD association, positive mediation effects were observed by *Lachnoclostridium* (13.3%), along with *Tyzzerella* and *Weissella*, and negative effects by *Ruminococcus* (−6.4%), *Eisenbergiella*, and *Adlercreutzia*.

**Table S6:**
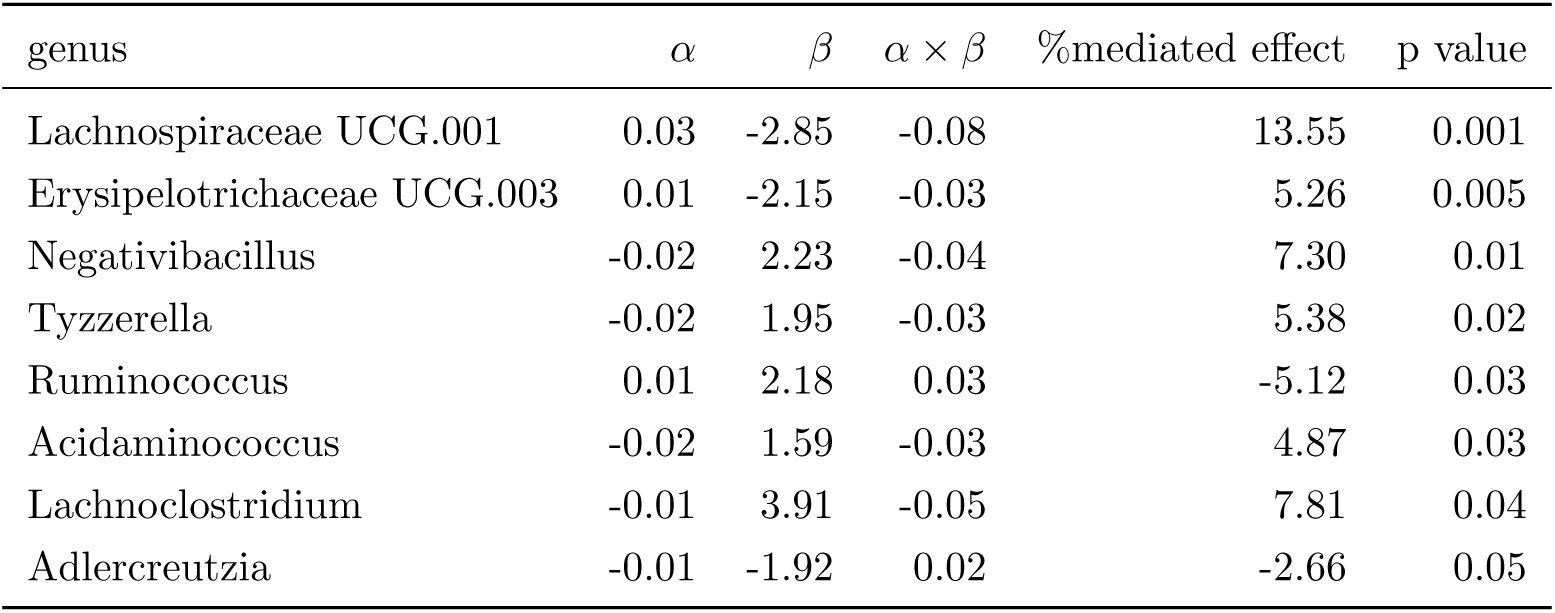
Genera mediating an association between *X*=HEI and *Y* =visceral fat. Total effect *γ* = −0.59.

**Table S7:** Genera mediating an association between *X*=HEI and *Y* =liver fat. Total effect *γ* = −0.05.

| genus | $\alpha$ | $\beta$ | $\alpha \times \beta$ | %mediated effect | p value |
| --- | --- | --- | --- | --- | --- |
| Lachnoclostridium | -0.01 | 0.69 | -0.008 | 14.29 | $7.5 \times 10^{-6}$ |
| Weissella | -0.01 | 0.30 | -0.002 | 3.92 | 0.01 |
| Anaerotruncus | -0.01 | -0.24 | 0.002 | -3.48 | 0.02 |
| Adlercreutzia | -0.01 | -0.20 | 0.002 | -3.15 | 0.02 |
| Lachnospiraceae UCG.001 | 0.03 | -0.17 | -0.005 | 8.80 | 0.02 |
| Bilophila | -0.01 | 0.18 | -0.002 | 2.93 | 0.03 |
| Clostridia UCG.014 | 0.02 | -0.14 | -0.002 | 3.82 | 0.03 |
| Oscillibacter | -0.01 | 0.28 | -0.003 | 5.96 | 0.03 |
| Collinsella | -0.02 | -0.15 | 0.002 | -4.36 | 0.04 |
| Acidaminococcus | -0.02 | 0.11 | -0.002 | 3.86 | 0.05 |

**Table S8:**
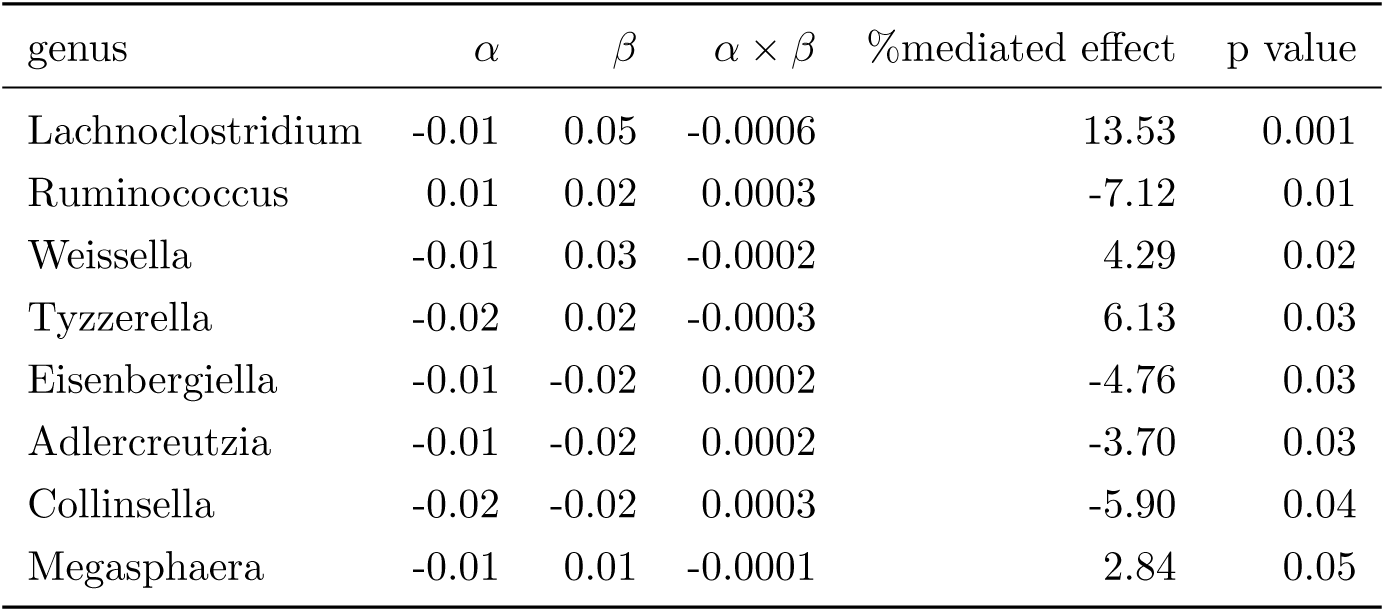
Genera mediating an association between *X*=HEI and *Y* =MASLD. Total effect *γ* = −0.004.

## Notes

### Competing Interest Statement

The authors have declared no competing interest.

